# TF protein of Sindbis virus antagonizes host type I interferon responses in a palmitoylation-dependent manner

**DOI:** 10.1101/2019.12.13.875781

**Authors:** Rogers KJ., Jones-Burrage S., Maury W., Mukhopadhyay S.

## Abstract

Sindbis virus (SINV) produces the small membrane protein TF from the 6K gene via a (−1) programmed ribosomal frameshifting. While several groups have shown that TF-deficient virus exhibits reduced virulence, mechanism(s) by which this occurs remain unknown. Here, we demonstrate a role for TF in antagonizing the host interferon response. Using wild-type and type 1 interferon receptor-deficient mice and primary cells derived from these animals, we show that TF controls the induction of the host interferon responses at early times during infection. Loss of TF production leads to elevated interferon and a concurrent reduction in viral loads with a loss of pathogenicity. Palmitoylation of TF has been shown to be important for particle assembly and morphology. We find that palmitoylation of TF also contributes to the ability of TF to antagonize host interferon responses as dysregulated palmitoylation of TF reduces virulence in a manner similar to loss of TF.

## Introduction

Alphaviruses are a genus of enveloped, positive sense RNA viruses comprised of a number of medically significant species. Old World alphaviruses such as chikungunya virus, Ross River virus, and Sindbis virus (SINV) cause debilitating arthralgias whereas New World alphaviruses such as Eastern, Western, and Venezuelan equine encephalitis viruses cause encephalitis and death (1, 2). Increased attention has been given to alphaviruses, which are spread by arthropod vectors, as broadening vector compatibility and global climate change alters the range and variety of vector species, resulting in increased outbreak frequency and severity (3-6).

Alphaviruses encode two open reading frames (ORFs) consisting of nonstructural (5’) or structural (3’) genes (7). Structural genes encode a large polyprotein consisting of capsid-E3-E2-6K-E1 which is subsequently proteolytically cleaved into individual proteins. It was recently discovered that within the gene encoding 6K there is a heptanucleotide slip site which causes a −1 ribosomal shift approximately 10-15% of the time (8, 9). This frameshift produces the truncated polyprotein capsid-E3-E2-TF, with the novel TF (<u>T</u>rans<u>F</u>rame) protein produced in place of 6K and E1 (9). Importantly the programmed frameshifting that produced TF is highly conserved amongst alphaviruses (10), with the heptanucleotide site, the downstream RNA regulatory elements, and the novel stop codon introduced by frameshifting found in all members of the genus (8, 10). Functionally, TF plays a role in viral assembly as loss of TF results in reduced virion release from cells and structural abnormalities (11). Furthermore, there is mounting evidence that TF plays a critical role in pathogenesis as viruses lacking TF are severely attenuated *in vivo* (12-15).

Post-translational modification of proteins is a common mechanism used by viruses to modify a protein’s function and cellular localization. Palmitoylation is the covalent attachment of a 16-carbon palmitoyl group primarily to cysteine residues which confers hydrophobic properties to proteins. Palmitoylated proteins are commonly targeted to cellular membranes (16). The palmitate group can be dynamically added and removed, thereby regulating the localization and function of proteins. Previous work has shown that wild type (WT) SINV TF is palmitoylated and exists in two states coined “basal” and “maximal”, both of which localize efficiently to the plasma membrane during viral assembly and release (11, 17). Mutating the cysteine residues in SINV that are palmitoylated causes defects in protein localization within the cell, TF incorporation into the virion, and virion morphology (11, 17). Palmitoylated TF is selectively incorporated into the released virion; however, the location of TF in the virion has not been identified in any structural studies. It is known that TF is present in lower, non-stoichiometric levels compared to the other structural proteins (capsid, E2, and E1) (18, 19). The low protein density due to the small size of TF and the low amounts and possible irregular distribution of the protein make it difficult to identify in reconstructions imposing icosahedral symmetry. Lastly, several C-terminal regions of the TF protein that regulate its own palmitoylation have been identified (17). Together these observations suggest that palmitoylation is a tightly regulated modification necessary for proper localization of TF and subsequent virion production.

RNA viruses are potent inducers of the host interferon (IFN) response and activate numerous pattern recognition pathways (20-22). Unsurprisingly, alphavirus replication and pathogenesis has been shown to be tightly restricted by IFN-α and -β production (IFN-I) (23, 24). Additional work has identified a number of interferon stimulated genes (ISGs) and pathways that control SINV virus infection (25-33). Mice that are type 1 interferon receptor-deficient (*Ifnar*^*-/-*^) are susceptible to an otherwise non-lethal SINV virus infection suggesting a critical role for this response *in vivo* (34). Due to the potency of the IFN-I response, RNA viruses encode a diverse array of mechanisms by which they antagonize both the production of and response to IFNs (20-22, 35). Alphaviruses are known to achieve this through the combined action of several proteins: nsP1 early in infection, and nsP2, nsp3, and capsid at later timepoints (36-45). Together these proteins indirectly antagonize IFN-I responses by shutting down host gene transcription and interfering with host transcription and protein synthesis.

In this study, we used both wild type (WT) and *Ifnar*^*-/-*^ mice to identify a novel role for TF as an antagonist of IFN-I expression. Furthermore, we show that palmitoylation is a critical determinant of the interferon antagonistic capacity of TF as *Ifnar*^*-/-*^ but not WT mice were susceptible to infection with palmitoylation mutants. Finally, wild-type TF diminishes early IFN-I responses, enhancing SINV replication, whereas IFN-I levels are elevated during infection with the mutant 6K only SINV (which produces the 6K but not the TF protein), resulting in production of low virus titers. Together, these findings provide compelling evidence that SINV TF controls IFN-I production. This report serves as the first observation that SINV virus antagonizes interferon production directly. We propose that this pathway works in parallel and complements the indirect mechanisms of innate immune control previously reported for alphaviruses.

## Methods

### Production of viruses

Infectious viruses were generated from cDNA clones as previously described (11). Linearized SINV AR86 cDNA plasmid or the described TF mutant derivatives were transcribed into infectious RNA *in vitro* via SP6 polymerase and a synthetic cap analog (New England Biolabs, Ipswich, MA). RNA was electroporated into baby hamster kidney (BHK) cells resuspended in phosphate-buffered saline (PBS) with 1500 Volts, 25 microFarads capacitance, and 200 ohms resistance in a 2 mm cuvette (BioRad). Cells were plated in MEM supplemented with non-essential amino acids, L-glutamine, and antibiotic-antimycotic solution (Gibco) in 10% fetal bovine serum and grown in a 37°C incubator in a controlled-humidity environment with 5% CO_2_ until significant cytopathic effect was observed at which point the media were harvested. Cellular debris were pelleted at 5000*xg* for 15 minutes. Infectious virus was measured in the supernatant as plaque-forming units (PFU) on BHK monolayers as described below. Our parental strain was AR86 with the 6K/TF coding region from SINV TE12. There are two amino acid differences between AR86 and TE12 in this region (VV to FI respectively at amino acid residues 29 and 30). This chimera approach was used because previous work on TF regulation was done in SINV TE12.

### Mice and ethics statement

This study was conducted in strict accordance with the Animal Welfare Act and the recommendations in the Guide for the Care and Use of Laboratory Animals of the National Institutes of Health (University of Iowa (UI) Institutional Assurance Number: #A3021-01). All animal procedures were approved by the UI Institutional Animal Care and Use Committee (IACUC) which oversees the administration of the IACUC protocols (Protocol #8011280). C57BL/6 *Ifnar*^*-/-*^ mice (a kind gift of Dr. John Harty, University of Iowa, Iowa City, IA) were bred and maintained at the University of Iowa. WT C57BL/6 mice were purchased from Jackson Labs (Stock # 000664). Both male and female WT mice 3-4 weeks of age were used, as well as male and female *Ifnar*^-/-^ mice 6-8 weeks of age.

### Western Blot

BHK cells were infected an MOI=5 and 24 hours later, media were removed and monolayers were washed with PBS prior to being lysed in SDS Lysis Buffer (10 mM Tris pH 7.4, 1% SDS, supplemented with fresh 1 mM PMSF and 1 ug/ml Leupeptin). Lysate concentrations were measured by using a bicinchoninic acid (BCA) assay (Pierce). Samples were mixed to a 1X final concentration with 2X Tricine sample buffer (450 mM Tris HCl [pH8.45], 12% [vol/vol] glycerol, 4% [wt/vol] SDS, 0.0025%[wt/vol] Coomassie blue G250, 0.0025% [wt/vol] phenol red) and heated for 2 minutes at 85°C or 95°C before gel electrophoresis on a precast 10-20% Tricine gel (Thermo Fisher Scientific) in Tricine running buffer (100 mM Tris base [pH 8.3], 100 mM Tricine, 0.1% SDS). Gels were transferred, probed using the appropriate antibody (capsid, E1/E2, TF, 6K), and imaged as described before (11). Purified virions were obtained by infecting BHK cells (MOI=5), collecting supernatant at 24hpi, and pelleted through a 27% sucrose cushion in a Ti 50.2 TI for 2:10 hrs at 100,000 *xg* at 15°C.

### Plaque assay

Serial dilutions of samples containing infectious virus were added to BHK monolayers for 1hr at room temperature under gentle agitation. Cells were overlaid with 1% low-melt agarose, 1× complete MEM, and 5% fetal bovine serum and plaques were detected at 48 hpi by formaldehyde fixation and crystal violet staining. Viral titers are reported as plaque forming units per mL.

### *In vivo* infections

Footpad infections were performed by restraining mice (TV-150 STD Braintree Scientific Inc.) and injecting 1000 PFU in 25µl of virus or PBS in the footpad of the right hindpaw. No anesthesia was used. Intracranial infections were performed by first anesthetizing mice with ketamine/xylazine mix (87.5 mg/kg ketamine, 12.5 mg/kg xylazine). The injection site was sterilized with 80% EtOH and infection was performed by gently inserting a 28G needle into the cranium and injecting 1000 PFU in 25µl of virus or PBS. Mice were monitored to ensure recovery from anesthesia and screened for adverse effects due to the injection. To assess viral load in the periphery, whole blood was isolated via facial vein puncture. Blood was allowed to clot for 30 minutes at RT and serum was isolated by centrifugation at 8000xg for 2 minutes. To harvest brains, mice were euthanized by rapid cervical dislocation and whole brains were removed by gross dissection before RNA was isolated as described below.

### Peritoneal cell isolation and infection

To isolate peritoneal cells, mice were euthanized by rapid cervical dislocation. The peritoneal cavity was immediately lavaged with 10 mL of cold RPMI1640 + 10%FBS + 1% Penicillin-Streptomycin solution. Cells were then washed and resuspended in fresh media. Forty-eight hours after plating, non-adherent cells were removed by washing with PBS to obtained enriched peritoneal macrophage cultures. Cells were subsequently infected with SINV (MOI=1) and RNA was isolated at various timepoints as described below.

### Growth kinetics

A confluent layer of BHK cells was infected with WT or mutant SINV (MOI=5) and incubated at room temperature for 1 hour to allow adherence to cells. Cells were subsequently washed 1x with PBS to remove unbound virus. Fresh media was added and cells were incubated at 37°C for the duration of the experiment. At various times post infection 200µl of media was removed for titering (described above) and replaced with an equal volume of fresh media.

### qRT-PCR

For RNA isolation from whole organs we used the gentleMACS tissue dissociation system with associated gentleMACS M tubes. Organs were placed in 1mL of TRIzol (Invitrogen) in M tubes and dissociated for 1 minute. RNA was isolated from tissue culture samples using TRIzol (Invitrogen.) All steps were performed according to the manufacturer’s specifications. A total of 1μg of RNA was subsequently converted to cDNA with the High Capacity cDNA Reverse Transcription Kit (#4368814) from Applied Biosystems. Quantitative PCR was performed using POWER SYBR Green Master Mix (#4367659) from Applied Biosystems utilizing a 7300 real time PCR machine from Applied Biosystems. 20ng of cDNA were used in each well. Primers used are as follows: HPRT forward 5’-GCG TCG TGA TTA GCG ATG ATG-3’, HPRT reverse 5’-CTC GAG CAA GTC TTT CAG TCC-3’, Sindbis forward 5’-CCA CTG GTC TCA ACA GTC AAA-3’, Sindbis reverse 5’-CTC CTT TCT CCA GGA CAT GAA C-3’, Interferon Beta forward 5’-AGC TCC AAG AAA GGA CGA ACA T-3’, and Interferon Beta reverse 5’-GCC CTG TAG GTG AGG TTG ATC T-3’

### Total protein quantitation

1×10^6^ peritoneal macrophages were infected with either WT or 6K-only SINV (MOI=3). At the indicated times post infection, cells were lysed with PBS + 1% SDS (1µl Pierce Universal Nuclease to reduce viscosity). 10µl of each sample was loaded onto a Mini-PROTEAN TGX Stain-Free Gel (BIO-RAD), transferred to a nitrocellulose membrane, and stained with REVERT Total Protein Stain Kit (LI-COR). Protein was quantified on a LI-COR ODYSSEY CLx. The blot was probed with a polyclonal antibody that detects SINV capsid protein to determine viral protein levels over time.

### Statistics

*In vitro* experiments were performed three times and are shown as compiled means with error expressed as standard error of the mean. Significance was determined by two tailed Student’s t-test (alpha=0.05). *In vivo* experiments were performed a minimum of two times. Significance was determined using Log-rank (Mantel-Cox) test. All statistics were calculated using GraphPad Prism software (GraphPad Software, Inc.).

## Results

### Characterization of mutant viruses

We have previously generated a series of TF mutants of SINV TE12. 6K only expresses 6K, but not TF. In this virus, the heptanucleotide slip site was mutated such that TF is not produced (8, 9, 11, 13, 14). We also have generated and characterized SINV encoding TF protein with altered palmitoylation patterns (called 4C and 9C) (10, 17). In these studies, we introduced these mutations into the neurovirulent AR86 strain of SINV (46) to assess the role of TF as a virulence factor. Our parental and mutant AR86 strains showed the same pattern of palmitoylation as observed with TE12 viruses (11, 17). Both the basal and maximal palmitoylated forms of TF were observed in WT SINV-infected cells, whereas SINV 6K only did not produce detectable levels of TF. The 4C mutant produced hyper palmitoylated TF and the 9C mutant produced non-palmitoylated TF, as evidenced by the molecular weight of TF from 4C or 9C infected cells running slower or faster than WT TF on SDS PAGE, respectively (Figure 1A). Furthermore, we determined, as seen before, only the palmitoylated forms of TF (present in WT and 4C) were incorporated into virions, but the non-palmitoylated form of TF present in 9C-infected cells was not. The replication kinetics of these four viruses were evaluated over the first 24 hours. While we observed minor growth defects of 4C, 9C, and 6K only viruses between 9-12 hours of infection compared to WT SINV, by 24 hours post infection titers were not significantly different, suggesting that altered palmitoylation or loss of TF does not significantly impact viral replication in BHK cells (Figure 1B).

**Figure 1:**
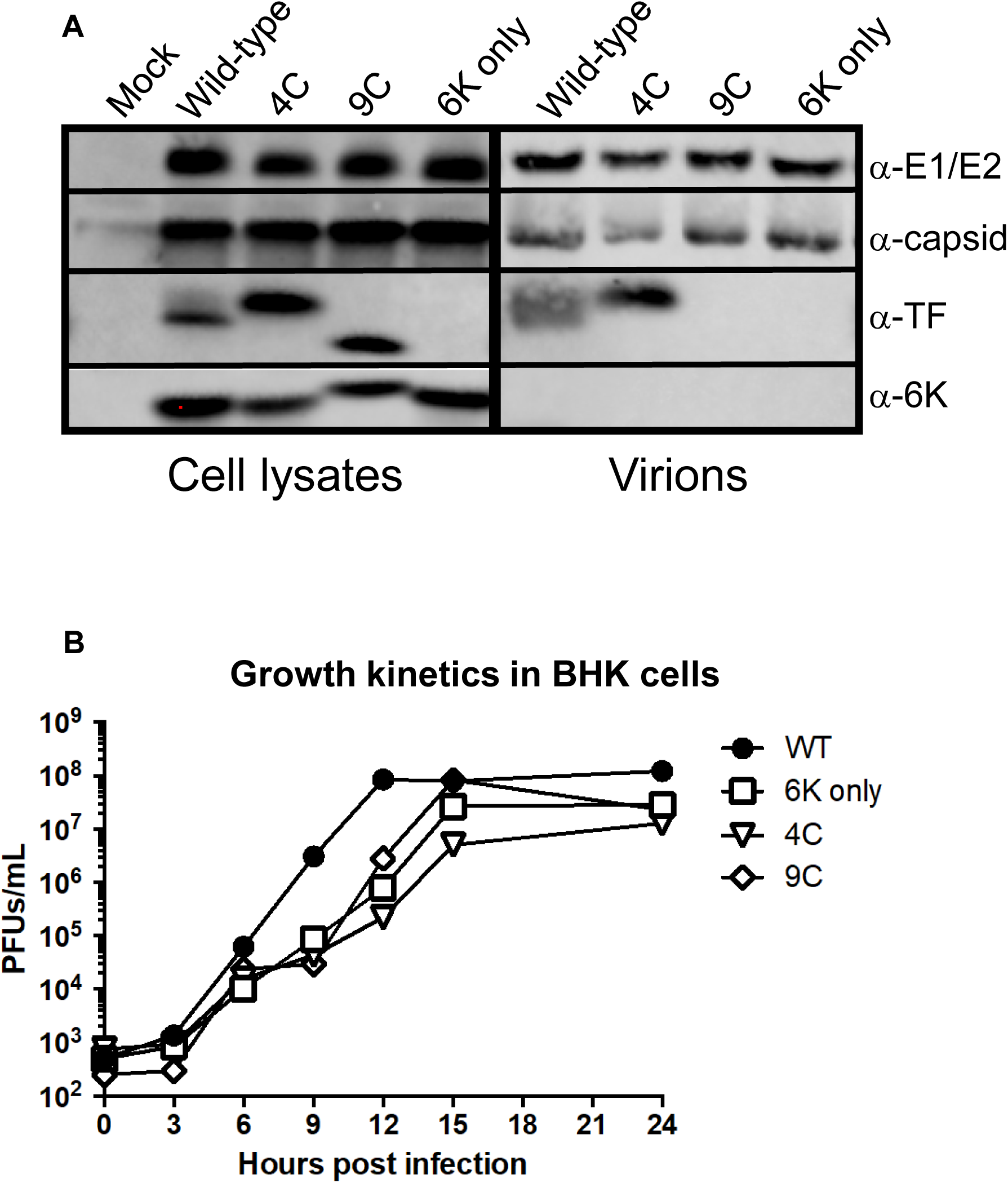
Characterization of viruses. **A)** BHK cells were infected with wild-type or TF mutant of SINV (AR86) (MOI=1). At 16 hours post infection, cells were lysed and lysates probed with antibodies specific for E1/E2, capsid, TF and 6K proteins. To probe components in purified virions, BHK cells were infected (MOI=5) and after 24 hours, media was collected, clarified, and run through a sucrose cushion. Pellets were resuspended and virion associated proteins were separated by SDS PAGE followed by western blotting. **B)** Growth kinetics were evaluated in BHK cells by plaque forming assays. Confluent monolayers of BHK cells were infected with the indicated virus (MOI=5) for 1 hour at room temperature. Cells were washed to remove unbound virus, media was refreshed, and cells were shifted to 37°C. At the indicated timepoints, 200µl of supernatant were removed and titered on BHKs to determine viral concentrations. Titers are reported as plaque forming units per milliliter. Each point represents the mean of two independent experiments.

### Palmitoylation of TF is a critical virulence determinant

Previous work demonstrated that the 6K only virus is attenuated in mice. Mice infected intracranially (IC) with the 6K only SINV survived while those infected with WT SINV succumbed to death (14); a similar result was seen with intranasally delivered Venezuelan equine encephalitis virus that lacked TF (13). To test the role of TF palmitoylation *in vivo*, we used a commonly used alphavirus model of footpad infection of young C57BL/6 mice (47-49) and the neurovirulent AR86 SINV strain. As infection progresses in this model, mice succumb to death. We found that ∼80% of mice succumbed to WT SINV infection, whereas all mice infected with 6K only virus survived, consistent with results from previous studies (13, 14). Interestingly, both the palmitoylation mutants, 4C and 9C, were severely attenuated (Figure 2A), just like the 6K only virus. Furthermore, infectious virus in the serum was significantly lower in mice infected with any of the mutants as compared to mice infected with WT SINV (Figure 2B). As we have previously found TF mutants to have altered virion morphology secondary to aberrant budding (11), we wanted to investigate *in vivo* trafficking defects as a possible explanation for this loss of virulence. We employed a second well-established model of infection, IC infection of young C57BL/6 mice (50, 51). Again, we found that all three mutant viruses had severely attenuated virulence in WT mice (Figure 2C). Furthermore, viral loads were significantly lower in the brains from mice infected with the mutant viruses as compared to WT (Figure 2D). Together with the growth kinetics in tissue culture these results suggest that loss of TF, or altered palmitoylation of the protein, directly attenuates SINV virus virulence.

**Figure 2:**
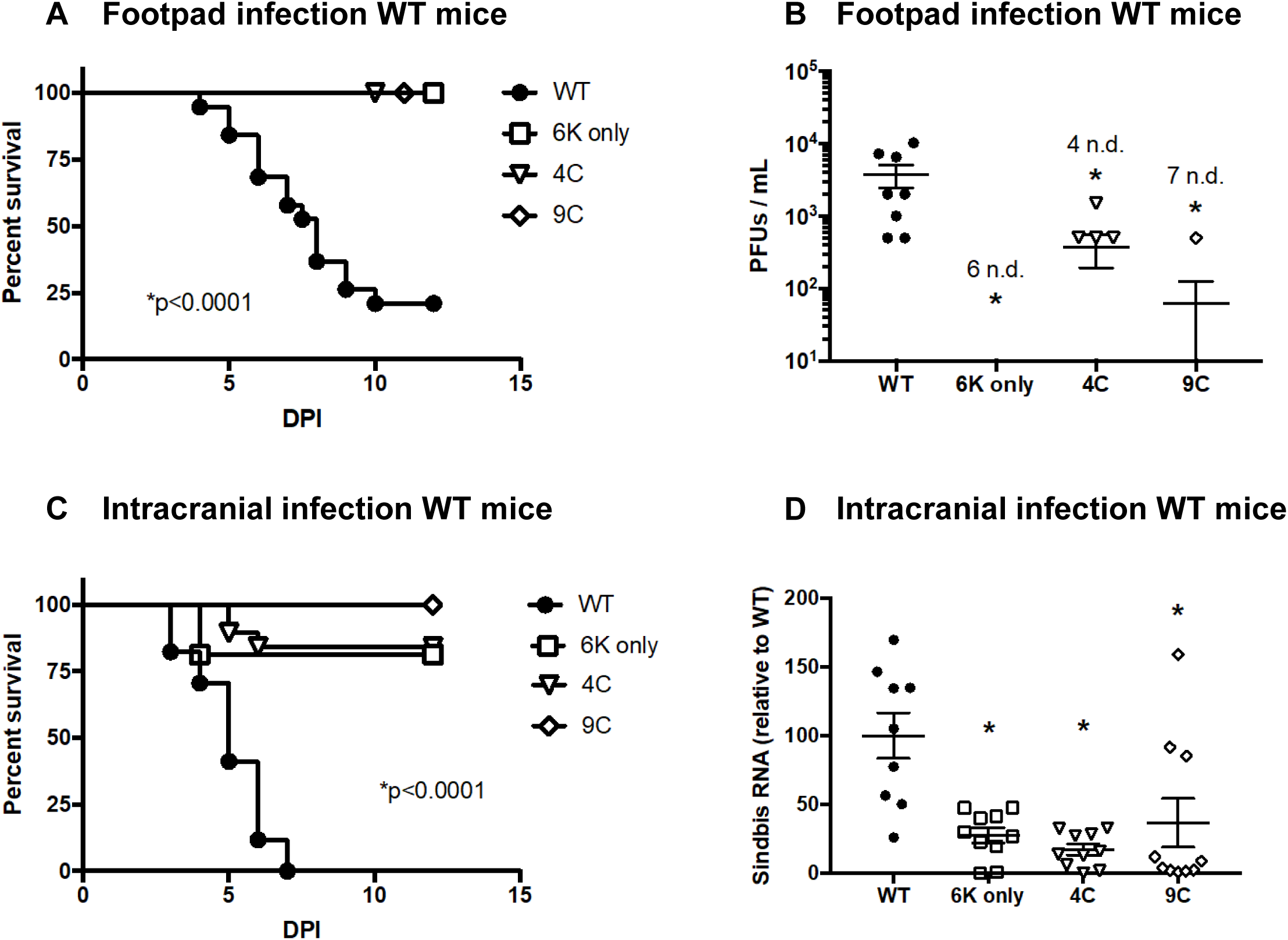
Virulence of SINV is dependent on expression and palmitoylation of TF. **A-B)** Male and female 3-4 week old C57BL/6 mice were infected with WT SINV (n=19), 6K only (n=17), 4C (n=15), or 9C (n=10) via footpad inoculation (1e3 PFU). Survival was monitored (A) and infectious virus in serum was determined by plaque assay at 36 hours post infection in a subset of mice (B). Experiment was performed 3 times. A value followed by n.d. indicates the number of samples in which no virus was detected. **C-D)** Male and female 3-4 week old C57BL/6 mice were infected with WT SINV (n=17), 6K only (n=16), 4C (n=19), or 9C (n=14) via intracranial inoculation (1e3 PFU). Survival was monitored (C) and viral load in brains was determined by qRT-PCR at 24 hours post infection in a subset of mice (D). This experiment was performed 3 times. Virus load determinations were normalized to the housekeeping gene, HPRT. Means +/- S.E.M. are shown for all viral quantitation along with data values for each mouse. Statistics for survival curves were performed using Log-rank (Mantel-Cox) test. Statistics for viral quantification were performed using Student’s t-test comparing mutant virus to WT.

As the TF mutant viruses are attenuated in WT mice but replicate well in BHK cells that lack IFN-I signaling, we hypothesized that WT SINV TF may play a role in antagonizing IFN-I responses dependent on its palmitoylation state. To determine if the presence of IFN-I in mice inhibits infection and virulence of TF mutants, we infected interferon α/β receptor knock out mice (*Ifnar*^*-/-*^) with WT and TF mutant viruses by footpad or I.C. We found that all of our viruses were virulent in *Ifnar*^*-/-*^ C57BL/6 mice with no significant difference in survival regardless of route of administration (Figure 3A and 3C). Furthermore, viral titer in serum from footpad infected mice (Figure 3B) or virus load in brains of IC infected mice (Figure 3D) did not differ significantly between WT and mutant viruses. Together these results suggest that TF, and its palmitoylation state, influence IFN-I-dependent immunity during SINV infection.

**Figure 3:**
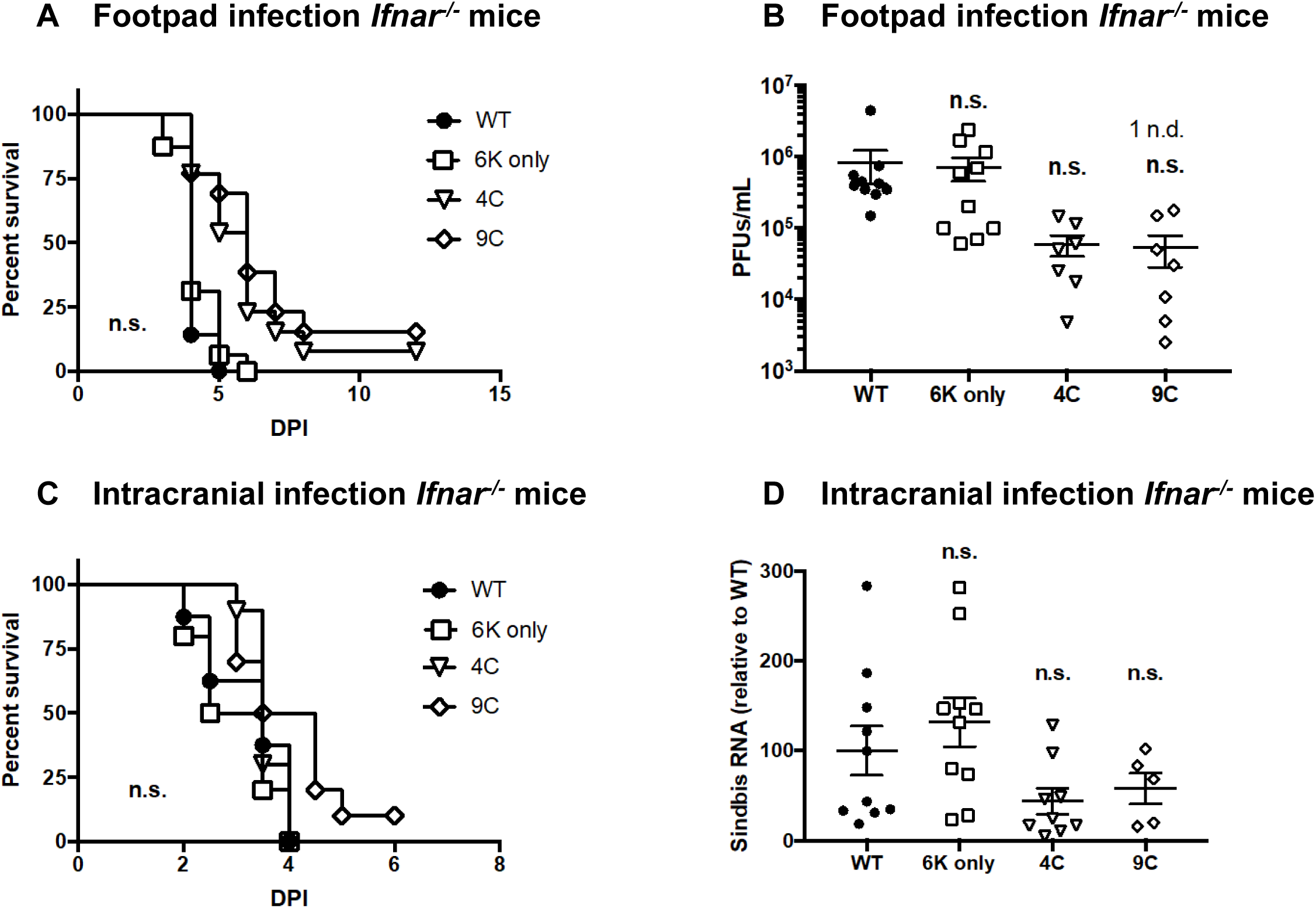
TF is not essential for virulence in type 1 interferon receptor deficient mice. **A-B)** Male and female 6-8 week old C57BL/6 *Ifnar*^*-/-*^ mice were infected with WT SINV (n=14), 6K only (n=16), 4C (n=13), or 9C (n=13) via footpad inoculation (1e3 PFU). Survival was monitored (A) and viral load in serum was determined by plaque assay at 36 hours of infection in a subset of mice (B). This experiment was performed 3 independent times. N.d. indicates samples in which no virus was detected. **C-D)** Male and female 6-8 week old C57BL/6 *Ifnar*^*-/-*^ mice were infected with WT SINV (n=10), 6K only (n=10), 4C (n=10), or 9C (n=10) via intracranial inoculation (1e3 PFU). Survival was monitored (A) and viral load in brains was determined by qRT-PCR at 24 hours of infection in a subset of mice (B). Virus load was normalized to the housekeeping gene, HPRT. Means +/- S.E.M. are shown for all viral quantitation along with data values for each mouse. The experiment was performed 2 times. Viral quantification data are expressed as mean +/- S.E.M. Statistics for survival curves were performed using Log-rank (Mantel-Cox) test. Statistics for viral quantification were performed using Student’s t-test comparing mutant virus to WT. ns = not significant.

### TF antagonizes host interferon responses

Given our *in vivo* observations, we directly tested whether TF in the context of WT SINV antagonizes the IFN-I responses. Resident peritoneal macrophages isolated from C57BL/6 mice were infected with WT or 6K only SINV *ex vivo*. Tissue macrophages were chosen as they are a critical component of the innate immune response and have been previously shown to be infected by SINV (22, 33, 47, 48). Cells were infected with virus and harvested over the first 24 h of infection. RNA was isolated and virus load and IFN-β expression were evaluated by qRT-PCR. Expression was normalized for the expression levels of the housekeeping gene HPRT. During these early times of infection, WT SINV produced significantly higher viral loads and lower IFN-β RNA synthesis than 6K only infection, implicating TF in controlling IFN-1 production (Figure 4A and 4B). We observed the same inverse relationship between WT SINV and IFN-β RNA *in vivo*. Infection of C57BL/6 mice IC with WT SINV resulted in higher virus loads and lower IFN-β expression in the first 24 hours of infection compared to 6K only virus (Figure 4C).

**Figure 4:**
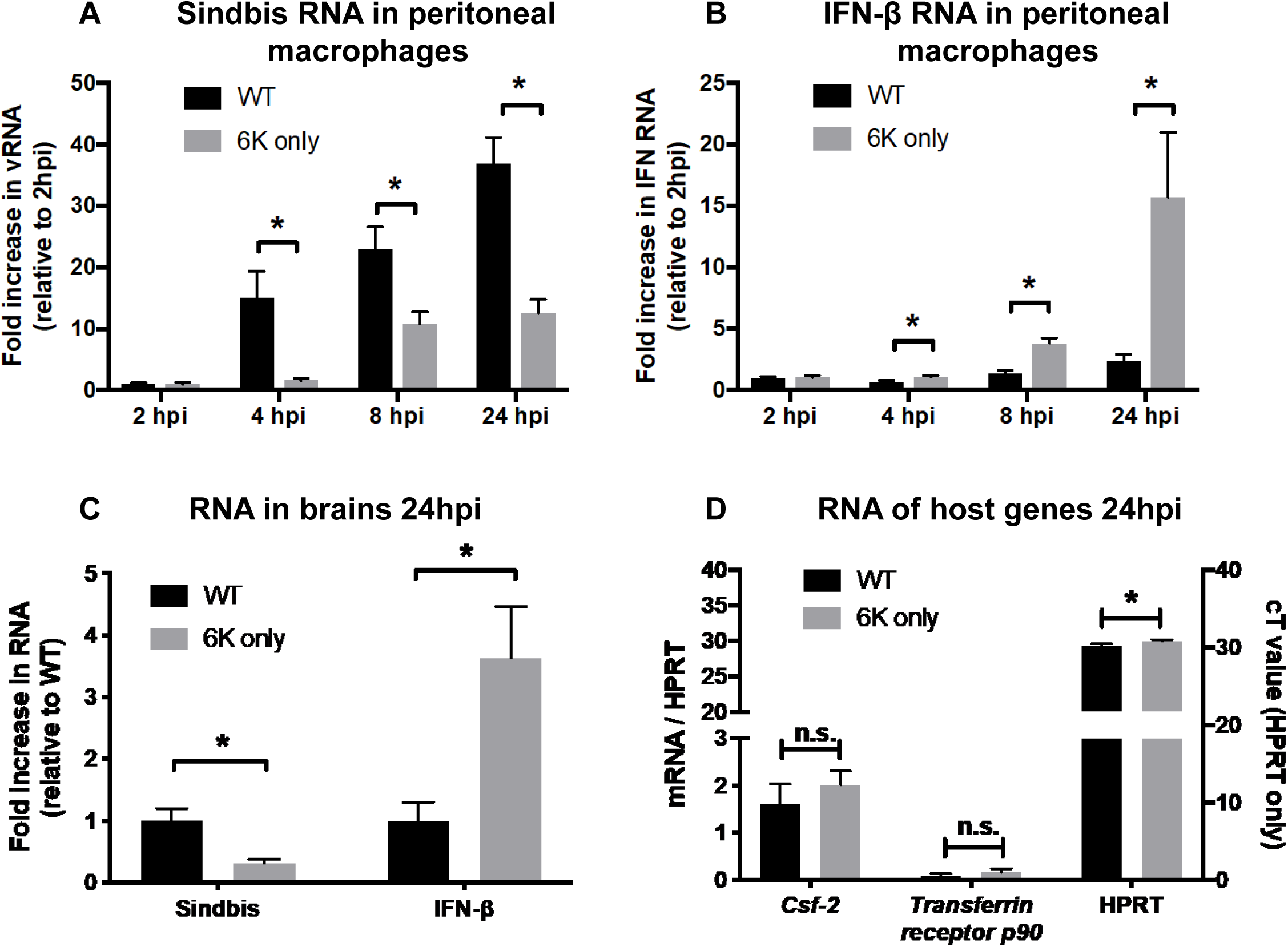
TF antagonizes type 1 interferon responses *in vitro* and *in vivo*. **A-B)** Peritoneal macrophages were harvested from female C57BL/6 mice and plated in tissue culture for 48 hours to allow adherence. Cultures were washed to remove non-adherent cells and subsequently infected with WT Sindbis or 6K only (MOI =1). RNA was harvested at the indicated timepoints and virus (A) and interferon beta (B) were quantified by qRT-PCR. The housekeeping gene HPRT was used to normalize expression. Data are expressed as mean +/- S.E.M. of fold changes relative to the 2hpi timepoint of WT infected cells to allow pooling of three independent experiments. **C)** Male and female 3-4 week old C57BL/6 mice were infected with WT Sindbis (n=10) or 6K only (n=10) via intracranial inoculation (1e3 PFU). 24 hours post infection mice were euthanized, brains were harvested, and RNA was isolated. Levels of Sindbis virus and interferon beta RNA were quantified by qRT-PCR. Data are expressed as mean +/- S.E.M. of fold changes relative to the WT infected mice to allow pooling of two independent experiments. **D)** Peritoneal macrophages were harvested from female C57BL/6 mice and plated in tissue culture for 48 hours to allow adherence. Cultures were washed to remove non-adherent cells and subsequently infected with WT or 6K only SINV (MOI =1). At 24hpi, RNA was isolated and samples were assessed for two host transcripts with high turnover rates. Data for *Csf-2* and *Transferrin receptor p90* are expressed relative to HPRT values on the left y-axis, and HPRT Ct values are shown on the right y-axis. Statistics were performed using Student’s t-test in all cases.

To determine if the changes in RNA levels were a consequence of a downstream effect of virus-induced host cell transcriptional shut off, we measured levels of the host genes *Csf-2* and *Transferrin receptor p90* in the WT SINV and 6K only infected samples. These mRNAs contain AU sequences in their 3’ untranslated region that cause RNA instability and rapid turnover of message, thereby serving as sensitive readouts of host transcriptional changes (52). At 24 h of SINV infection, we did not observe differences in either their raw Ct values (data not shown) or expression of these genes relative to the housekeeping gene HPRT (Figure 4D). Interestingly, the absolute level of HPRT was slightly, but statistically significantly, lower (higher Ct value) in the 6K only compared to the WT SINV infection, indicating slightly more host cell HPRT mRNA was present in cells infected with WT virus. When comparing the levels of host and viral proteins in WT and 6K only-infected cells, there were no differences over the first 24 hours of infection (data not shown). Together these findings indicate that the decreased level of IFN-β in WT SINV infections compared to 6K only infections is a consequence of the TF protein and not because of differences in host cell transcription or translation regulation. In total, these *ex vivo* and *in vivo* experiments provide evidence that TF blocks the initiation of IFN-β responses and that this is a specific effect rather than due to overall virus suppression of host cell transcription and/or translation. These results provide insight into a novel role for TF protein and a mechanistic understanding of TF as a virulence factor.

## Discussion

Previous studies have demonstrated that the TF protein of a number of different alphaviruses serves as a virulence factor (13, 14); however, the mechanism responsible for TF-induced virulence has not been identified. Here, we provide evidence in mice and primary macrophages that SINV TF acts as an IFN-I antagonist. TF was not needed for SINV virulence in *Ifnar*^*-/-*^ mice regardless of whether virus was delivered peripherally or into the brain. However, TF expression was required for virulence in WT mice, independent of the route of delivery. Consistent with a role for TF in controlling IFN-I production, wild-type SINV infection of macrophages resulted in low IFN-β RNA levels and concomitantly high virus loads. 6K only SINV-infected mice had higher IFN-I production and low virus titers. Together our data support a model for TF antagonizing IFN-I production.

Our studies are the first to implicate alphavirus TF as an antagonist of IFN-I synthesis. Viruses from other RNA virus families are well established to interfere with both IFN-I synthesis and downstream IFN-I signaling (35). Further, our data supports the contention that TF may specifically inhibit IFN-I synthesis since differences in host cell transcription and/or translation were not detected. The antagonistic activity of TF may complement the activity of other alphavirus proteins. Nsp1, nsp2, and capsid have been previously shown to either control host innate immune responses downstream of IFN-I production and/or directly inhibit host cell transcription and translation (36-45). Utilizing multi-prong mechanisms for escaping from or interfering with host defenses is a strategy used by many viruses. Here, we have identified another piece of the puzzle used by alphaviruses and identified a new role for the viral protein TF.

We previously found that palmitoylation of TF was critical for its localization to the plasma membrane, TF incorporation into progeny virus, and virus morphology (11). SINV mutants that lack palmitoylation (9C) or are hyper palmitoylated (4C) were not virulent in wild-type mice. However, *Ifnar*^*-/-*^ mice were susceptible to infection with palmitoylation mutants, providing evidence that the palmitoylation status of TF controls the interferon antagonist capacity of TF. Taken together, the proper palmitoylation state of TF maybe be required for inhibiting IFN production and regulating correct particle assembly and stability. While both mechanisms may indeed be at play, in mice the impact of TF on interferon antagonism is most notable, as the palmitoylation mutants and WT virus have indistinguishable virulence in *Ifnar*^*-/-*^ mice. Further, this highlights the multi-functional nature of TF; our data suggests that any growth defect, due to mis-palmitoylation that results impaired virus assembly/release, appears independent of the anti-interferon function.

These studies provide evidence that TF interferes with IFN-I synthesis rather than globally blocking host cell transcription and/or translation. However, how TF interferes with IFN-production has yet to be examine. TF from the viral particle and/or newly synthesized in the target cells may bind and inhibit the activity of one or more host proteins involved in vRNA sensing, similar to hepatitis C (53), Zika (54), and other viruses (35). Aberrant TF palmitoylation may directly interfere with TF/host protein interactions. Alternatively, as the absence of palmitoylation or mis-palmitoylation alters SINV TF localization (11) during assembly, TF mutants may not appropriately traffic to cellular site(s) that are responsible for evading vRNA sensing pathways. Another possibility is that TF antagonism of IFN-I synthesis may be due to an indirect mechanism whereby TF binds and masks vRNA, thereby reducing the detection of incoming viral RNA present in the cell. Under this scenario, vRNA in our mutant viruses is more exposed and thus detected by the host vRNA sensors. Our preliminary analysis of TF stoichiometry in the particle and the surface properties of TF (including being a transmembrane protein and having few basic residues) makes this latter mechanism less likely. Future studies are planned to determine the mechanism by which TF blocks IFN-I synthesis.

## Acknowledgments

This work was funded by a grant from the Institute for Advanced Study at Indiana University (SM) and NIH grants AI139902 and AI134733 (WM).

